# FRMPD2: a novel GluN2A-interacting scaffold protein in synaptic excitatory transmission

**DOI:** 10.1101/403360

**Authors:** Xi Lu, Haiqing Zhang, Xin Tian, Xuefeng Wang

## Abstract

The scaffold proteins FRMPD2 is localized at the basolateral membranes of polarized epithelial cells and associated with tight junction formation. But the expression and function of FRMPD2 in vivo remain unknown. Here, we found that postsynaptic FRMPD2 effectively alters the neuronal excitability by regulating NMDARs-component excitatory synaptic transmission in adult hippocampus. FRMPD2 anchors the NMDAR subunit GluN2A at the synapses without changing the total NMDAR expression. By using of Co-IP, Pull Down and SPR, the second PDZ (PDZ2) domain of FRMPD2 direct and binds to the C-terminus of GluN2A. Our study discover that a novel scaffold protein FRMPD2 regulates synaptic excitatory in a PDZ2 domain-dependent manner in adult hippocampus.

## Introduction

Scaffold proteins contain several modular protein interaction domains within their structure, and they bind to various interacting proteins, such as ion channels, neurotransmitter receptors, adhesion molecule, microtubule, filamentous actin (F-actin)(Anzai et al., 2002; Durand et al., 2012; Elias and Nicoll, 2007; Hruska-Hageman et al., 2002; Kim and Sheng, 2004; Sala et al., 2001; Schultz et al., 1998). They strongly contribute to the organization of synaptic signalling complexes, and the clustering of surface receptors and ion channels, and they participate in the dynamic trafficking of vesicle transport and coordinate cytoskeletal dynamics(Feng and Zhang, 2009). FRMPD2 (also named Gm626 in mice) is a novel scaffold protein that consists of an N-terminal KIND domain, a Ferm (band 4.1, ezrin, radixin, and moesin) domain and three C-terminal PDZ (PSD-95/Discs-large/ZO1) domains(Lipinski et al., 2012; Stenzel et al., 2009). FRMPD2 is reported to localize in the basolateral membranes of polarized epithelial cells and is associated with tight junction formation and immune response(Lipinski et al., 2012; Stenzel et al., 2009). However, there are few studies investigating FRMPD2 in vivo and the expression and function of FRMPD2 in nervous system remain unknown.

Inspired by multiple domains in its structure, FRMPD2 is predicted to play a critical role in neuronal excitation. PDZ domain-containing proteins are involved in neuronal excitatory by trafficking neurotransmitter receptors to plasma membrane(Kim and Sheng, 2004). Thus, FRMPD2 may be implicated in the regulation of neuronal excitation in a PDZ- dependent manner.

In this paper, we first reported that postsynaptic scaffold protein FRMPD2 activated neuronal excitability in adult hippocampus by mediating the N-methyl-d-aspartic acid (NMDA) receptors-component excitatory synaptic transmission. FRMPD2 anchored the GluN2A subunit of NMDA receptors (NMDARs) to the synapses by directly binding the second PDZ (PDZ2) domain to the C-terminus of GluN2A. Our study shed light on a novel scaffold protein FRMPD2 mediate synaptic excitatory via directly anchoring synapses GluN2A subunits.

## Results

### 1. Expression patterns of the FRMPD2 protein

The FRMPD2 protein was expressed in a restricted manner in some organs including the brain, eyes, testes and lungs (Fig 1A), and was enriched in a more widespread manner in various brain regions, including the cortex, hippocampus and cerebellum, in addition to the ependymal cells lining the ventricles (Fig 1B, C).

**Fig 1.**
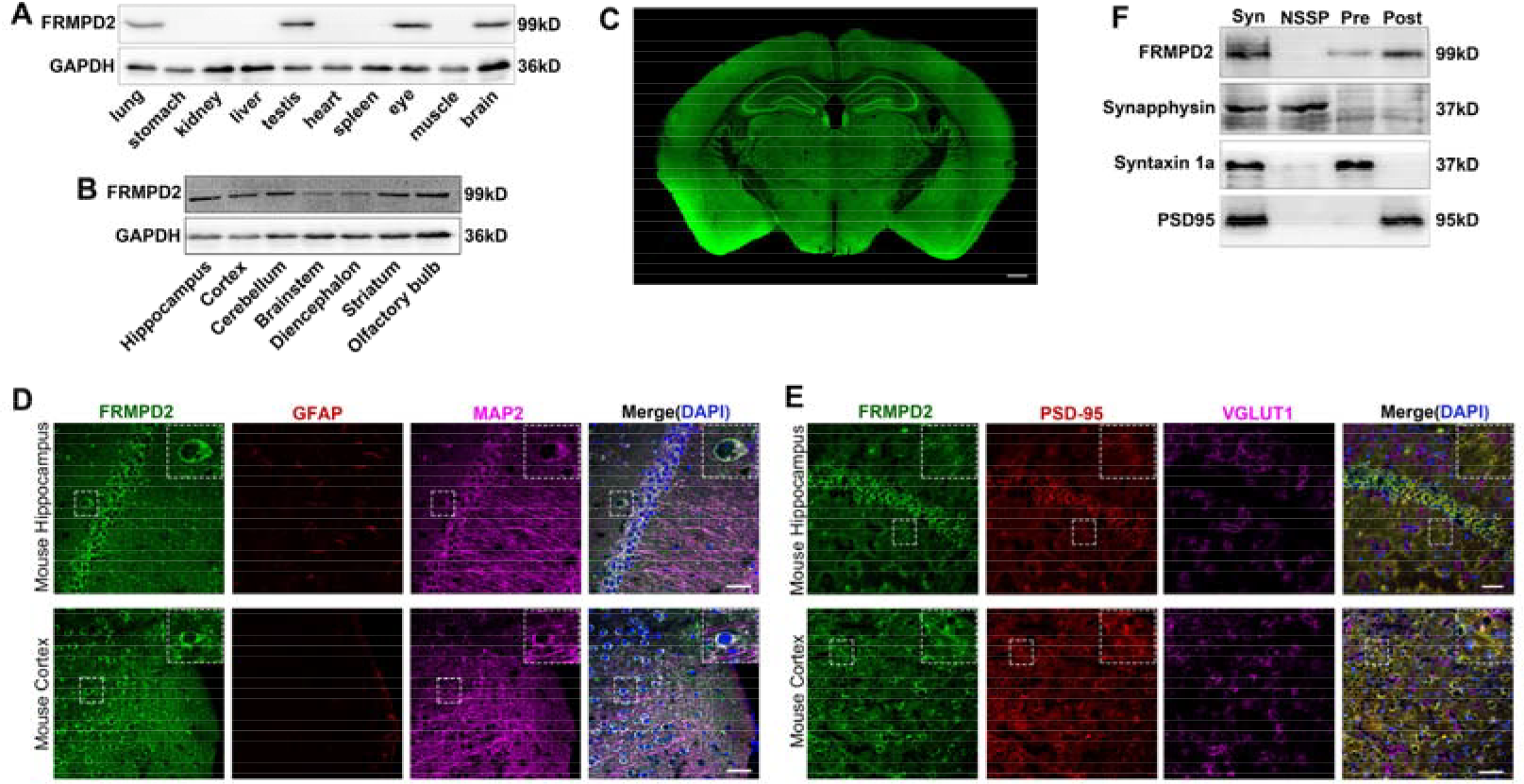
Expression patterns of the FRMPD2 protein. A, FRMPD2 protein is located in the mouse brain, eyes, testes and lungs. B, The brain-specific expression of the FRMPD2 protein was revealed in adult mouse tissue homogenates. C, The FRMPD2 protein has a widespread distribution pattern in adult mouse brain slices. Bar, 500 μm. D, FRMPD2 was present in MAP2-positive neuronal dendrites but not in GFAP-positive glial cells as determined by immunofluorescence staining. Bar, 40 μm. E, FRMPD2 co-localized with the postsynaptic marker PSD95 but not with the presynaptic marker VGLUT1 in the mouse hippocampus and cortex. Bar, 40 μm. F, Immunoblot of the synapse fraction (Syn) showing that FRMPD2 was preferentially distributed in the postsynaptic fraction (Post) rather than in the presynaptic (Pre) or non-synaptic synaptosomal proteins (NSSP) fraction. Anti-synapphysin, anti-syntaxin 1a and anti-PSD95 were used as markers for ultrasynaptic fraction to verify that the purity of the preparation was higher than 90%. The blots were representative of 4-6 analyses.

When immunofluorescence staining was performed in mouse cortex and the hippocampus Cornu Ammonis 1 (CA1) region, FRMPD2 was localized in microtubule-associated protein 2 (MAP2)-positive neuronal dendrites and another major scaffold protein, PSD95 (postsynaptic marker) (Fig 1D, E). Synaptosomes in the hippocampus were isolated by ultracentrifugation, and the FRMPD2 protein was detected in the postsynaptic density (PSD) fraction, but not in the presynaptic fraction or non-synaptic synaptosomal fraction (Fig 1F).

### 2. Effects of FRMPD2 on NMDA receptor-mediated excitatory synaptic transmission

PDZ domain-containing proteins are reported to be involved in neuronal excitatory (Kim and Sheng, 2004). We tested that whether postsynaptic FRMPD2 containing three PDZ domains affects neuronal excitatory. Whole cell patch-clamp electrophysiology was performed in CA1 pyramidal neurons in mouse hippocampal slices at 14 days after lentivirus vector (LV) injection of mice that was used to knockdown or overexpress FRMPD2. LV-encoding green fluorescent protein (GFP) was detected in the mouse hippocampus (Fig 2A). At 2 and 6 weeks post-LV injection, the FRMPD2 protein levels of the hippocampus in FRMPD2 knockdown mice (shFRMPD2) was almost 40 percent less than in scramble controls (shFRMPD2-Scr). FRMPD2 protein levels were significantly higher in shFRMPD2-Res mice that were injected with shRNA plus overexpression LV in the hippocampus than in FRMPD2 knockdown mice. The FRMPD2 protein in FRMPD2 overexpression mice (FRMPD2-OE) was 1.9-2.1 times higher than in negative controls (FRMPD2-OE NC) (Fig 2B). These findings suggest that FRMPD2 protein levels in the hippocampus were effectively modulated by LV injection.

**Fig 2.**
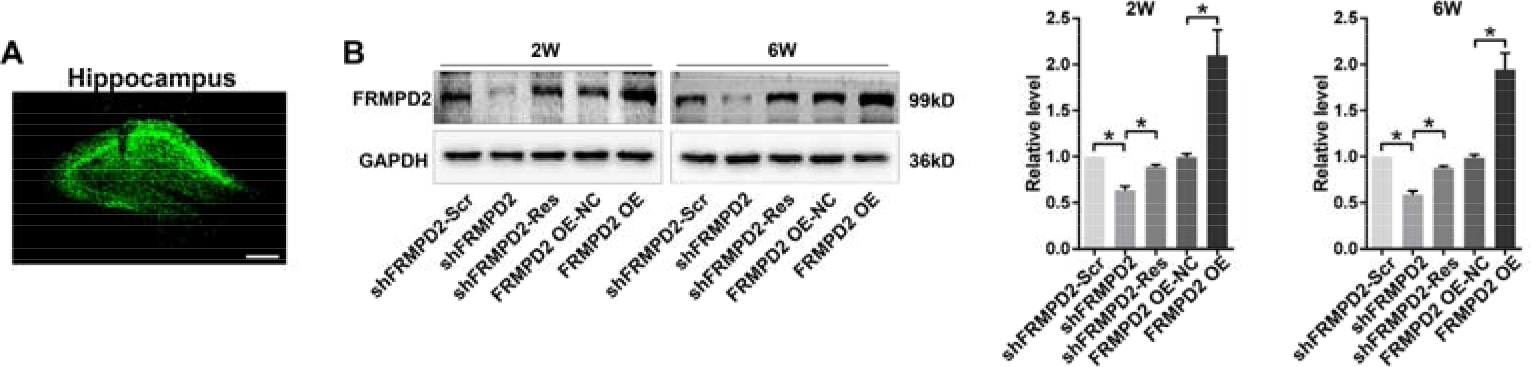
Alteration of FRMPD2 protein levels in the hippocampus by LV injection. A, Lentiviral vector (LV)-encoding green fluorescent protein (GFP) was detected in the mouse hippocampus following the injection of LV. Bar, 250 μm. B, Hippocampal FRMPD2 protein expression levels were detected in FRMPD2 knockdown or overexpression mice on the 2 and 6 weeks after LV injection. Data were shown as mean±SD and analyzed by Student’s t test, n=5 mice, *p< 0.05.

We firstly analyzed the effect of FRMPD2 on spontaneous action potential (AP) in CA1 pyramidal neurons. The frequency of spontaneous AP in FRMPD2 knockdown mice reduced by 71 percent compared with the scramble control. AP frequency of FRMPD2 overexpression mice was 2.28 times faster than in the negative control (Fig 3A). To further explore the contribution of glutamate excitatory receptors and/or inhibitory synaptic inputs contribute to the increase in neuronal hyperactivity induced by FRMPD2, □-amino-3-hydroxy-5-methyl-4-isoxazole propionic acid (AMPA) receptor (AMPARs) component or NMDAR component mEPSCs, and miniature inhibitory postsynaptic currents (mIPSCs) of pyramidal neurons in the CA1 region of the hippocampus were recorded. The two types of ionotropic glutamate receptors, AMPARs and NMDARs, present at postsynaptic membrane of excitatory synapses mediate almost all synaptic transmission in the brain(Bredt and Nicoll, 2003; Valtschanoff and Weinberg, 2001). There was no significant difference in AMPAR mEPSCs (Fig 3B) or mIPSCs (Fig 3C) after FRMPD2 knockdown or overexpression. However, the mEPSC amplitude, but not the frequency, of NMDAR mEPSCs was decreased in FRMPD2 knockdown mice compared with scramble controls, and increased in FRMPD2 overexpression mice compared with negative controls (Fig 4A).

**Fig 3.**
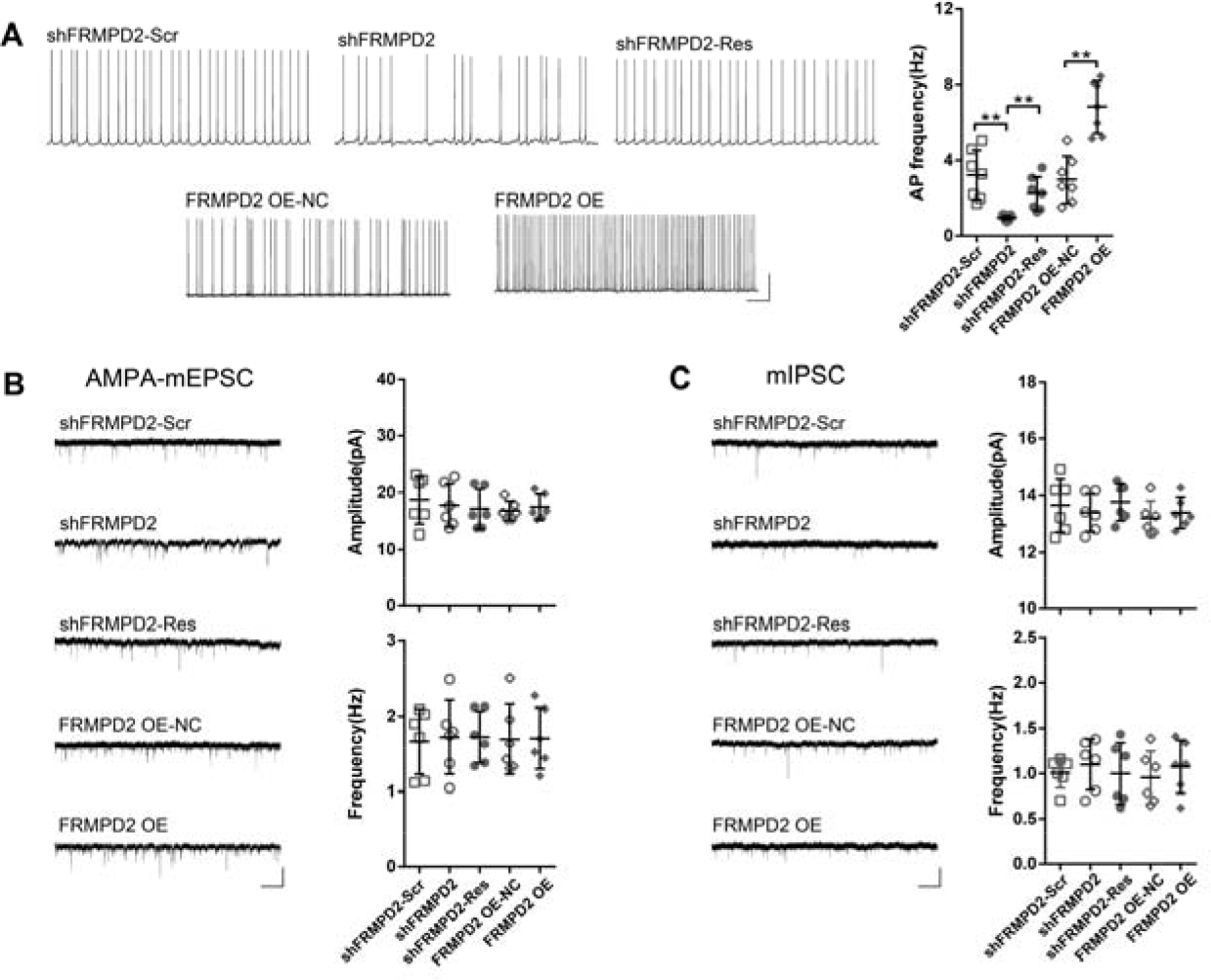
Effects of FRMPD2 on electrophysiology of CA1 neurons in mouse hippocampal brain slices. A, The frequency of spontaneous AP was compared after FRMPD2 knockdown or overexpression. Bar, 1 s, 25 mV. B, C, FRMPD2 had no significant effect on the amplitude or frequency of AMPAR mEPSCs (B) and mIPSCs (C). Bar, 1 s, 20 pA. Data were shown as mean±SD and analyzed by Student’s *t* test, n=6-7 neurons of 3 mice, **p< 0.01.

**Fig 4.**
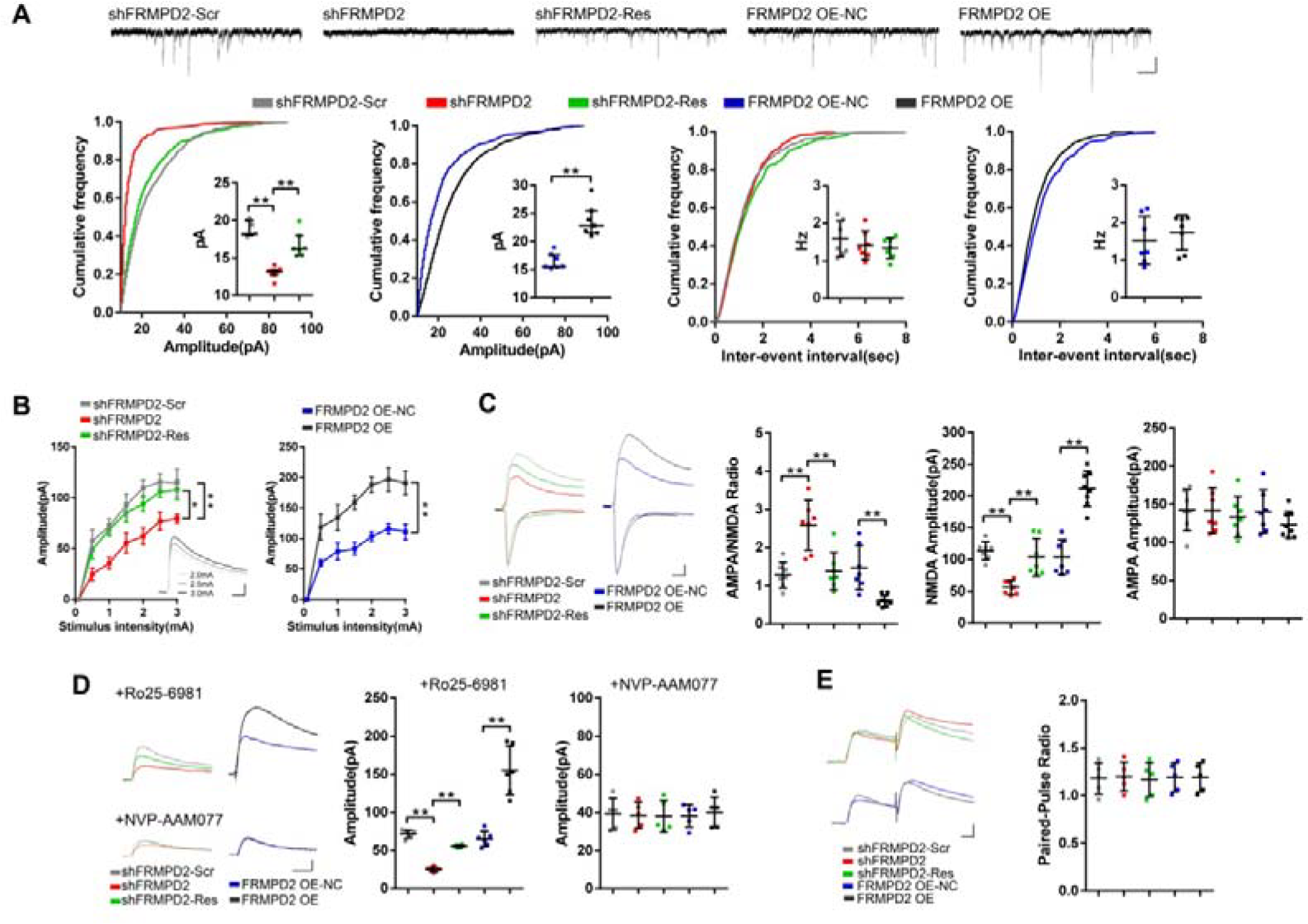
Effects of FRMPD2 on NMDA receptor-mediated excitatory synaptic transmission. A, Quantitative analysis of amplitude and frequency of NMDAR mEPSCs in CA1 pyramidal cells of hippocampus after FRMPD2 knockdown or overexpression. Bar, 1 s, 20 pA. B, The input-output curves of NMDAR eEPSCs were recorded with gradual increasing intensities. Bar, 25 ms, 50 pA. C, Amplitude of NMDAR, AMPAR component eEPSCs and AMPA/NMDA ratio were compared. Bar, 25 ms, 25 pA. D, Quantitative analysis of the amplitude of GluN2A- and GluN2B-sensitive eEPSCs. Bar, 25 ms, 25 pA. E, Knocking down or overexpressing FRMPD2 did not alter the paired-pulse ratio (PPR) values. Bar, 25 ms, 50 pA. Data of were presented as the mean±SD or median with interquartile range. Data of A, C, D, E, were analyzed by Student’s *t* test, data of B was analyzed by repeated measures ANOVA, n=5-7 neurons from 3 mice. **p<0.01. *p<0.05.

To verify the changes of mEPSCs, NMDAR-mediated evoked EPSCs (eEPSCs) were measured. Firstly, the input-output curve of NMDAR eEPSCs (Fig 4B) verified that FRMPD2 enhanced the postsynaptic efficacy of NMDAR responses (Shen et al., 2010). As shown in Fig 4C, the amplitudes of NMDAR currents were significantly lower in the FRMPD2 knockdown than in the scramble controls. There was no significant difference in evoked AMPAR currents between the FRMPD2 knockdown and overexpression groups, in line with data of AMPAR component mEPSCs. In particular, to determine whether GluN2A and/or GluN2B component currents contributed to the observed changes in NMDAR eEPSCs, selective antagonist for GluN2A (NVP-AAM077) and GluN2B (Ro25-6981) were used (Fig 4D). The amplitude of GluN2A-sensitive currents of the FRMPD2 knockdowns was decreased by 65 percent compared with the scramble groups, and amplitude of GluN2A currents of overexpressing FRMPD2 mice was 2.38 times higher than of the negative controls. Interesting, the GluN2B component-mediated currents were not changed after FRMPD2 knockdown or overexpression. These results suggested that FRMPD2 strengthens GluN2A -mediated NMDARs synaptic transmission.

Because presynaptic release at postsynaptic glutamate receptors also contribute to synaptic excitatory transmission, we analyzed the paired-pulse ratio (PPR), a reliable index of the contribution of presynaptic vesicles release to excitatory postsynaptic currents(Zucker and Regehr, 2002). There was no significant difference in PPR values between the FRMPD2 knockdown and overexpression results and those in the corresponding control groups (Fig 4E), indicating that presynaptic vesicles release is not responsible for the FRMPD2-mediated changes in mEPSCs

### 3. Effects of FRMPD2 anchoring GluN2A at synapses

To correlate electrophysiological changes with biochemical alterations in synaptic NMDAR, we performed immunoblotting using hippocampal synaptosomal and total lysates. As shown in Fig 5A, the synaptic GluN2A and GluN2B were significantly fewer in the FRMPD2 knockdown mice than in the scramble mice, and higher in the FRMPD2 overexpression mice than in the negative controls. The total protein levels of GluN2A, GluN2B and GluN1 were not affected by FRMPD2. Immunofluorescence staining was also used to test for co-localization between FRMPD2 and GluN2A at postsynaptic membranes, which were marked by PSD-95 in hippocampal CA1 areas (Fig 5B).

**Fig 5.**
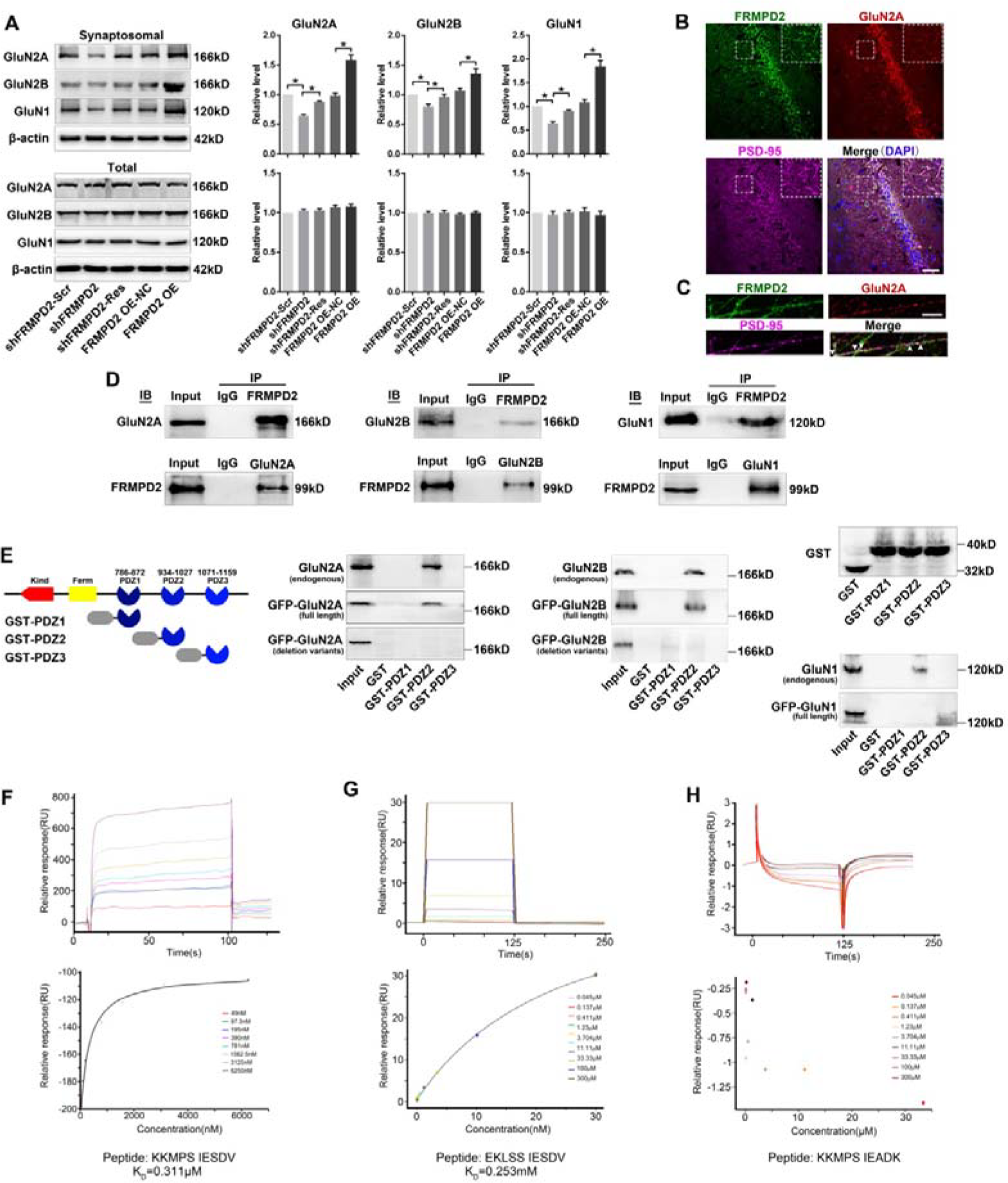
Effects of FRMPD2 anchoring GluN2A at synapses. A, Quantitative analysis of GluN2A, GluN2B and GluN1 expressions of total hippocampus tissue and synaptic homogenates after FRMPD2 knockdown or overexpression. The data were shown as the mean±SD and analyzed by Student’s *t* test, *p< 0.05. B, Immunofluorescence staining in the mouse hippocampus CA1 area showed that FRMPD2 co-localizes with GluN2A at the postsynaptic density, which was marked by PSD95. Bar, 40 μm. C, FRMPD2 and GluN2A, GluN2B and GluN1 were co-immunoprecipitated from mouse hippocampal tissues. D, GST pull-down assay showing that the GST-PDZ2 fusion protein but not the PDZ1 or PDZ3 fusion proteins pulled down endogenous GluN2A from mouse hippocampal tissues, and exogenous full-length GFP-GluN2A from HEK293 cells but did not pull down the deletion variant GFP-GluN2A from HEK293 cells transfected with GFP-GluN2A that lacked ten amino acids at the C-terminus. E-G, The ability of the isolated FRMPD2 PDZ2 domain to bind to the GluN2A and GluN2B C-terminus was measured using SPR. Peptides sequences of C-termini of wild type GluN2A (E), wild type GluN2B (F) and mutant GluN2A (G) were used in this study. Sensorgrams were shown to illustrate the binding of increasing concentrations of the PDZ2 domain to peptides and steady-state equilibrium responses from the sensorgrams were shown plotted against the injected PDZ2 concentrations. K_D_: dissociation constant.

The extreme C-terminus of the NMDAR subunits provides a mechanism for trafficking NMDAR subunits in postsynaptic membrane by interacting with PDZ proteins (Bard et al., 2010; Kim and Sheng, 2004). We performed co-immunoprecipitation (Co-IP) and GST pull-down assays to detect whether the three PDZ domains of FRMPD2 interact with NMDAR subunits. In vivo, co-immunoprecipitation of FRMPD2 with GluN1, GluN2A and GluN2B were detected in the lysates of mice hippocampus (Fig 5C). A GST fusion protein bearing the PDZ2 domain but not the first PDZ or third PDZ domain pulled down endogenous GluN1, GluN2A and GluN2B from mice hippocampus in vivo. The full-length GluN1, GluN2A and GluN2B proteins tagging with GFP were generated in translation experiments in vitro, GST-PDZ2 pulled down full-length GluN2A and GluN2B, but not GluN1 (Fig 5D). However, the GST-PDZ2 fusion protein did not pull down GluN2A proteins from the cells transfected with a form of GFP-GluN2A that lacked a set of ten C-terminal amino acids (Fig 5D).

The binding abilities of FRMPD2-isolated PDZ2 domain to the GluN2A and GluN2B C-terminus was measured using surface plasmon resonance (SPR) analyses. The binding capacity of the FRMPD2 PDZ2 domain to bind to the wild type GluN2A C-terminus (KKMPSIESDV) and wild type GluN2B C-terminus (EKLSSIESDV) yielded a K_D_ (dissociation constant) value of 0.31μM and 0.25mM, respectively (Fig 5E,F), whereas no binding was observed between the mutant GluN2A C terminus (KKMPSIEADK) (Fig 5G) and the FRMPD2 PDZ2. These results supported the direct interaction of C terminus of GluN2A to PDZ2 of FRMPD2.

To further explore potential sites for binding between GluN2A and FRMPD2, we first obtained the crystal X-ray structure of the PDZ2 domain(Fig 6A) of the mouse homologue of FRMPD2 (PDB ID: 5ZDS) (Table 1). The crystal structure was solved by PHENIX with the template PDZ2 domain structure of tyrosine phosphatase PTP-BL (PDB ID: 1VJ6) (Fig 6B) with 63% identity(Adams et al., 2010; Gianni et al., 2006). In the template structure (Fig 6C), there was a peptide that is a C-terminal (HSGSYLVTSV) of tumor suppressor protein APC, binding to β2 strand and α2 helix. The residues V(0) and T(−2) of the peptide formed critical hydrophobic interactions with residues L25, I27, L85, V29, and V65 of the template PDZ2 structure(Gianni et al., 2006). These residues were highly conserved in the FRMPD2 PDZ2 crystal structure, as shown by the alignment (Fig 6D).

**Table 1.**
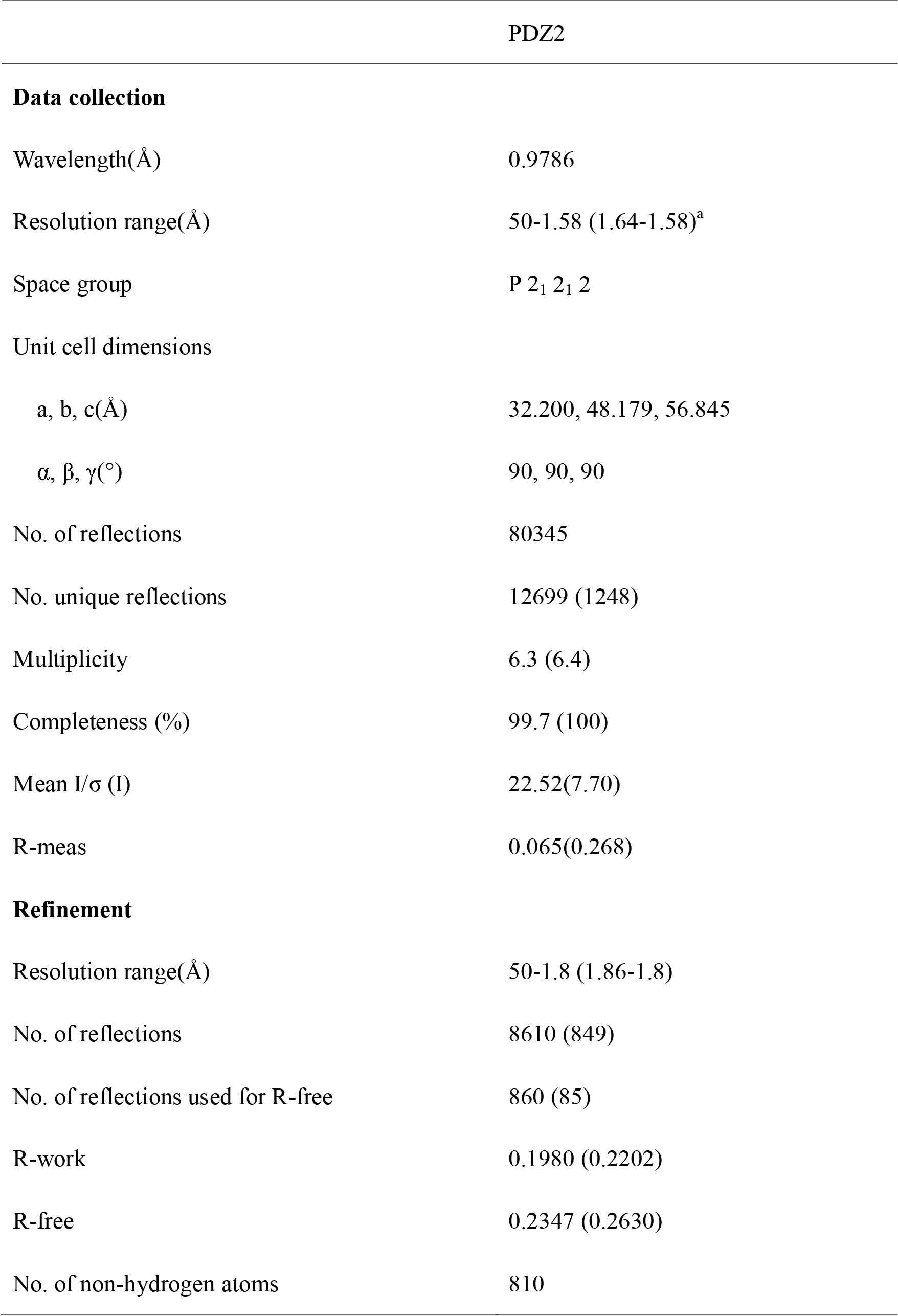
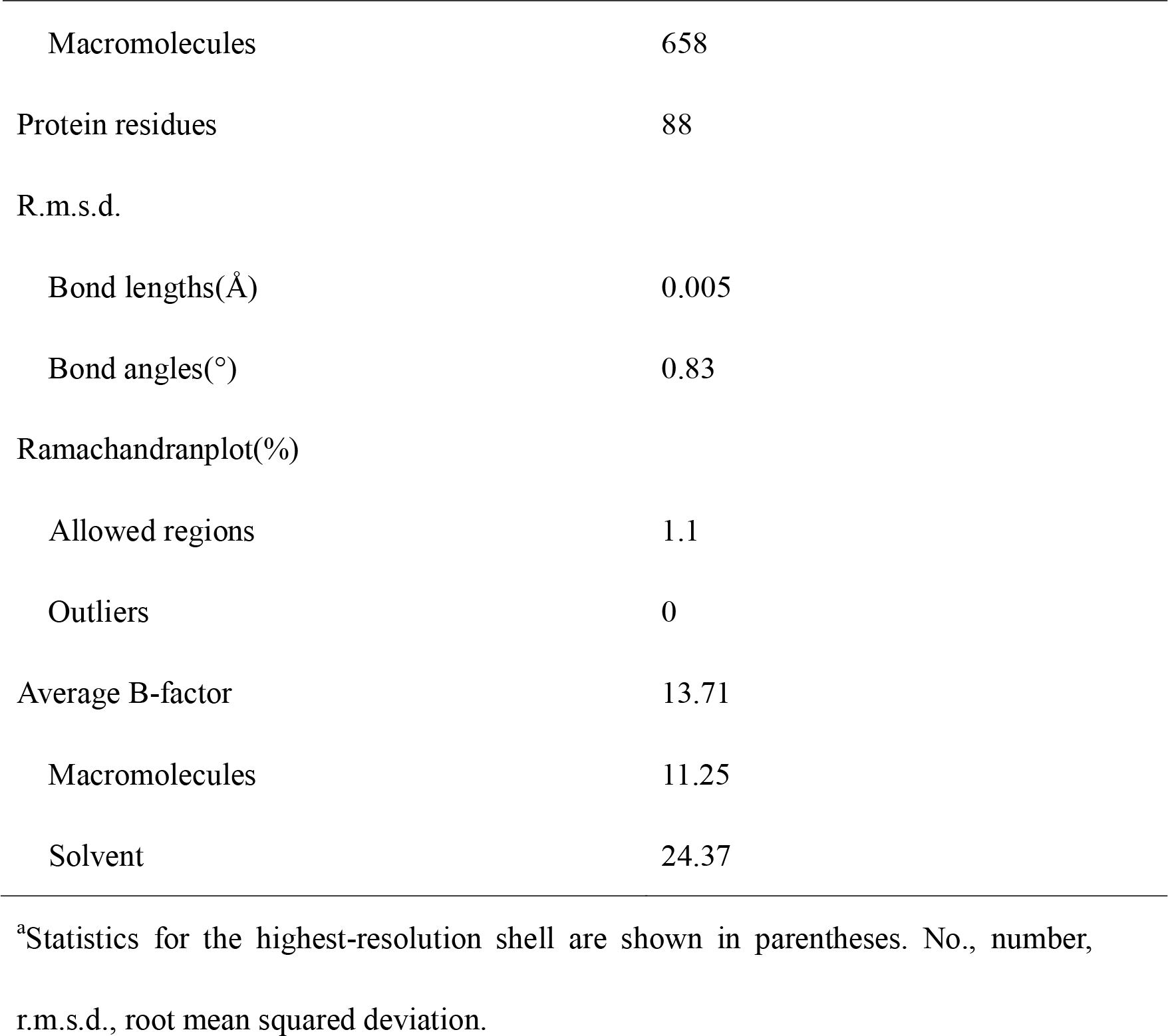
Data collection and refinement statistics of PDZ2 of FRMPD2.

|  | PDZ2 |
| --- | --- |
| **Data collection** |  |
| Wavelength(Å) | 0.9786 |
| Resolution range(Å) | 50-1.58 (1.64-1.58)<sup>a</sup> |
| Space group | P 2<sub>1</sub> 2<sub>1</sub> 2 |
| Unit cell dimensions |  |
| a, b, c(Å) | 32.200, 48.179, 56.845 |
| $\alpha$, $\beta$, $\gamma$(°) | 90, 90, 90 |
| No. of reflections | 80345 |
| No. unique reflections | 12699 (1248) |
| Multiplicity | 6.3 (6.4) |
| Completeness (%) | 99.7 (100) |
| Mean I/$\sigma$ (I) | 22.52(7.70) |
| R-meas | 0.065(0.268) |
| **Refinement** |  |
| Resolution range(Å) | 50-1.8 (1.86-1.8) |
| No. of reflections | 8610 (849) |
| No. of reflections used for R-free | 860 (85) |
| R-work | 0.1980 (0.2202) |
| R-free | 0.2347 (0.2630) |
| No. of non-hydrogen atoms | 810 |
| Macromolecules | 658 |
| Protein residues | 88 |
| R.m.s.d. |  |
| Bond lengths(Å) | 0.005 |
| Bond angles(°) | 0.83 |
| Ramachandranplot(%) |  |
| Allowed regions | 1.1 |
| Outliers | 0 |
| Average B-factor | 13.71 |
| Macromolecules | 11.25 |
| Solvent | 24.37 |

<sup>a</sup>Statistics for the highest-resolution shell are shown in parentheses. No., number, r.m.s.d., root mean squared deviation.

**Fig 6.**
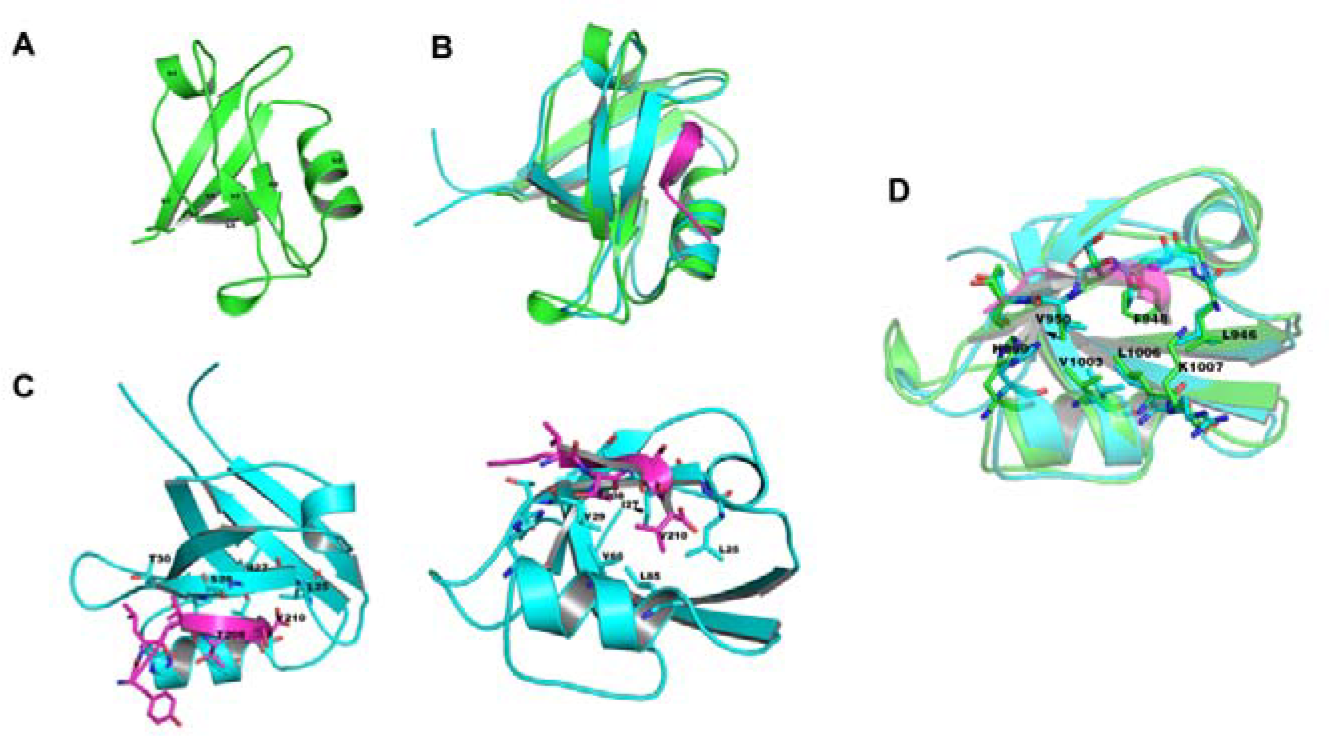
Crystal structure of FRMPD2 PDZ2 domain. A, The crystal structure of a compact globular fold in the ligand-free FRMPD2 PDZ2 domain (PDB ID: 5ZDS) was shown. B, The crystal structure of the FRMPD2 PDZ2 domain (green) was shown aligned to that of the PDZ2-peptide template structure (cyan and magenta) (PDB ID: 1VJ6). C, A template of PDZ2-peptide (cyan and magenta) interaction (PDB ID: 1VJ6). D, The binding pocket alignment between FRMPD2 PDZ2 crystal structure (green) and PDZ2 template (cyan).

The results shown so far reveal that FRMPD2 affects NMDAR-mediated excitatory postsynaptic currents by anchoring GluN2A subunits at postsynaptic membranes. This activity is achieved by an interaction of the FRMPD2 PDZ2 domain to the GluN2A C-terminus.

## Discussion

In our report, we identified a novel scaffold protein FRMPD2, and explored its physiological contribution to the modulation of excitatory synaptic electrophysiological function.

FRMPD2 was essential for NMDA-component, but not AMPA-component excitatory transmission in adult hippocampus. It is seems that the binding of glutamate receptors by FRMPD2 appears to change with glutamate receptor-specific manner(Beique et al., 2006; Elias et al., 2008). Our results showed that synaptic FRMPD2 mediated neurotransmission by trafficking GluN2A subunits on synapses in adult hippocampus. NMDA receptors are composed of obligatory GluN1 subunit co-assembled with GluN2A or GluN2B subunit (GluN1/GluN2A or GluN1/GluN2B). The identity of the GluN2A and/or GluN2B subunits determines the biophysical and pharmacological properties of NMDARs (Bard et al., 2010; Sultan et al., 2015). Our results of Co-IP, GST-pull-down and SPR assays identified the second PDZ domain of FRMPD2 as a novel binding partner of GluN2A and GluN2B *in vivo* and *in vitro*. Although the FRMPD2 PDZ2 did not directly bind to GluN1, the obligatory GluN1 subunit of GluN1/2A complex was, expectedly, detectable by Co-IP of FRMPD2. Meanwhile, the GluN1 protein of synaptosome was changed after FRMPD2 knockdown or overexpression, which was accompanied by the alteration of GluN2A subunits. In addition, the binding capacity of PDZ2 to GluN2A subunit was 1000 times higher than that to GluN2B subunit. It is probably that the binding of PDZ2 to GluN2B is flexible and instant. In addition, the GluN2B-containing NMDARs are located at extrasynaptic sites and involved in neurotoxicity in adult hippocampal neurons, GluN2A-containing NMDARs mainly resided at synaptic sites and involved in neurotransmission (Parsons and Raymond, 2014).

In summary, the results presented here suggest a critical role of novel scaffold protein FRMPD2 in regulating NMDA receptors anchoring, which is critical for regulation of glutamatergic synapses transmission. The dysregulation of this scaffold protein contributed to the impaired excitatory conductance.

## Methods and materials

### DNA constructs

To construct the following GST fusion proteins, the PDZ domain of mouse FRMPD2 (Uniprot: A0A140LI67) was subcloned into pGEX4T-1 with a GST tag: GST-PDZ1 (amino acids 786-872) and GST-PDZ2 (amino acids 934-1027), GST-PDZ3 (amino acids 1071-1159).

The lentivirus vector (LV) constructs were manufactured by Genechem Co. Biotechnology (Shanghai City, China). The mouse FRMPD2 (NCBI Gene: 268729) short hairpin RNA (shRNA) knockdown construct targeted 5’-3’ CTGAATCTTTCTCTTGCAT and was subcloned it into the LV GV248 (GeneChem, China), which contained green fluorescent protein (GFP). A scramble shRNA (5’-3’ CTTTTCTAACGTGTCGACT) was used as a negative control for the FRMPD2 shRNA. The complete mouse cDNA sequence for FRMPD2 (XM_017316244) (FRMPD2 overexpression, FRMPD2 OE) and the GFP sequence were inserted into the LV GV358 (GeneChem, China).

### Antibodies

The following antibodies were used: FRMPD2 (rabbit, Sigma;rabbit, Origene), Flag(mouse, Sigma), GST(rabbit, Invitrogen), GFP (rabbit, Invitrogen), GluN2A(rabbit, Proteintech), GluN2B(mouse, Proteintech; rabbit, Abcam), GluN1(rabbit, Abcam), MAP2(guinea pig, SYSY), GFAP(mouse, Sigma), PSD-95(guinea pig, SYSY; mouse, Millipore), VGLUT1(guinea pig, SYSY), synaptophysin(rabbit, Abcam), syntaxin 1a(rabbit, Abcam).

### Cell line cultures and transfection

HEK293T and SH-SY5Y cells were maintained in DMEM (Gibco) supplemented with 10% fetal bovine serum (Gibco), 100 U/mL penicillin and 100 μg/mL streptomycin (Gibco) at 37°C in 5% CO2. The cells were transfected using Lipofectamine 2000 (Invitrogen)(Klein et al., 2017). At 4 h after transfection, the cells were rinsed with warm PBS and incubated in culture medium containing 10% fetal bovine serum.

### Intrahippocampal injection of LV

Adult C57/BL6 male mice were obtained from the Experimental Animal Center of Chongqing Medical University. Intrahippocampal injection of LV was performed using a stereotaxic apparatus (Stoelting Co. Ltd Wood Dale, IL, USA). Briefly, the mice were anesthetized with pentobarbital (80mg/kg, intraperitoneal) and placed on stereotaxic apparatus. The reference points were bregma for the anterior-posterior axis, the midline for the medial-lateral axis and the dura mater for the dorsal-ventral axis with a tooth bar. LV particles were injected bilaterally into the dorsal hippocampus through a glass pipette (0.2 μl/min) attached to a glass microsyringe(Shangguan et al., 2017). The mice were allowed to recover for 2 weeks after the vector injections.

### Patch clamp recordings

Brain slices were prepared as previously reported(Chen et al., 2006; Kim et al., 2001). Briefly, C57/BL6 mice were anesthetized with pentobarbital. Brain slices (300 μm) were prepared with a Leica (Germany) VP1200S Vibratome and incubated in artificial cerebral spinal fluid (ACSF) (119 mM NaCl, 26 mM NaHCO_3_, 2.5 mM KCl, 1 mM MgCl_2_, 1.25 mM NaH_2_PO_4_, 2 mM CaCl_2_ and 25 mM glucose, pH 7.4, 310 mOsm) bubbled with 5% CO_2_ and 95% O_2_ for at least 1 h at room temperature before recording. Whole-cell recording was performed as described previously(Liu et al., 2005). Glass microelectrodes (Sutter, USA) were shaped by a pipette puller (P-97, Sutter, USA) to a final resistance of 3–5 MΩ when the pipette was filled with internal solution. A multi-clamp 700B amplifier (Axon, USA) was used for the recordings. Signals were sampled at 10 kHz and filtered at 2 kHz. A stable baseline was obtained for at least 5 min prior to recording and data were discarded when the access resistance (15–20 MΩ) was changed by 20% at the end of recording. Mini Analysis 6.0.1 (Synaptosoft) and Clamp Fit 10.3 software (Axon) were used to analyze the recorded data.

For action potential (AP) recording, glass pipettes were filled with the following internal solution: 17.5 mM KCl, 0.5 mM EGTA, 122.5 Mm K-gluconate, 10 mM HEPES, and 4 mM ATP, pH adjusted to 7.2 with KOH [20]. The internal solution used to record excitatory post-synaptic currents (EPSCs) contained: 17.5 mM CsCl, 10 mM HEPES, 4 mM ATP, 0.5 mM EGTA, 132.5 mM Cs-gluconate, and 5 mM QX-314. The internal solution used to record inhibitory post-synaptic currents (IPSCs) contained: 100 mM CsCl, 1 mM MgCl_2_, 1 mM EGTA, 30 mM N-methyl-D-glucamine, 10 mM HEPES, 5 mM MgATP 0.5 mM Na_2_GTP and 12 mM phosphocreatine.

Spontaneous activity was assessed by observing the neuron under current clamp for 1 min. Tetrodotoxin (TTX, 1 μM), bicuculline (10 μM), 6,7-dinitroquinoxaline-2,3(1H,4H)-dione (DNQX, 20 μM) and D-serine (10 μM) were added to Mg-free-ACSF to record NMDAR mEPSCs that held at −60 mV. TTX (1 μM), bicuculline (10 μM) and amino-5-phosphonopentanoate (AP-V, 40 μM) were added to ACSF to record AMPA mEPSCs. TTX(1 μM), DNQX(20 μM) and APV(40 μM) were added to ACSF to record mIPSCs that held at −70 mV (Shangguan et al., 2017).

Evoked EPSCs (eEPSCs) currents were generated with a 40μs pulse (0.1 Hz) from a stimulated isolation unit controlled by an AMPI generator (Master-8, USA). A bipolar stimulation electrode (FHC) was located ~100 μm rostral to the recording electrode in the same layer(Chen et al., 2008; Tang et al., 2015). The eEPSCs at −70 mV holding potential in the present of PTX (100 μM) was considered as the AMPAR-mediated current. Neurons were then held at +40 mV and the amplitude of the eEPSCs at 50 ms post-stimulus was interpreted as the NMDAR-mediated current(Etherton et al., 2009; Etherton et al., 2011). Input-output curve of NMDAR eEPSCs were recorded in response to a series of stimulation intensities (pA). Paired-pulse ratio (PPR) recordings were voltage-clamped at +40 mV and obtained using a paired-pulse protocol of two stimuli at an inter-pulse interval of 50 ms. PPR values were defined as the ratio of the second peak amplitude to the first peak amplitude. The highly selective GluN2B subunit antagonist Ro25-6981 (Ro, 0.5μM) was applied to assess GluN2A-sensitive eEPSCs, and the selective GluN2A antagonist NVP-AAM077(NVP, 0.4μM) was added to record GluN2B component eEPSCs(Bartlett et al., 2007).

### Western blotting

Samples boiled with 4X sample buffer at 95°C for 5 min, and then separated on 8-10% SDS-PAGE gels and transferred onto polyvinylidene difluoride membranes (Millipore). The target proteins were immunoblotted with primary antibodies overnight at 4°C and then incubated with HRP-conjugated secondary antibodies. The blots were imaged and quantified using a Fusion Imaging System. The quantitative densitometric values of the proteins were normalized to that of GAPDH or beta-actin.

### Immunofluorescence

For animal tissues, brain were post-fixed at 4°C for 24 h. Floating slices (50 mm in thickness) were permeabilized with 0.5% Triton X-100 and blocked in goat serum. The slices were then incubated with primary antibodies overnight at 4°C followed by incubation with secondary fluorescent conjugated antibodies at room temperature for 2 h. Images were captured using laser-scanning confocal microscopy (Nikon A1+R Microsystems) on an Olympus IX 70 inverted microscope (Olympus) equipped with a Fluoview FVX confocal scanning head. Areas of overlap and fluorescence intensity were analyzed using Image Pro Plus 6.0.

### Sub-synaptic fractionation

Sub-synaptic fractionation was performed as previously described(Phillips et al., 2001). To obtain synaptosomes, adult mouse hippocampus were homogenized in solution buffer (0.32 M sucrose, 0.1 mM CaCl_2_, 1 mM MgCl_2_, 0.1 mM PMSF) and brought to a final sucrose concentration of 1.25 M. Protease inhibitors were used in all purification steps. The homogenate was overlaid with 1.0 M sucrose and centrifuged (100,000 g, 3 h, 4°C). Synaptosomes were collected at the 1 M/1.25 M interface.

For sub-synaptic fractionation, synaptosomes were sequentially extracted with 1% Triton X-100 first at pH 6.0 and then at pH 8.0. Synaptosomes (described above) were diluted 1:10 with 0.1 mM CaCl_2_ to a final concentration of 20 mM Tris, pH 6.0 and 1% Triton X-100. After the solution was incubated for 30 min and then centrifuged 40,000 g for 30 min at 4°C. The supernatant (presynaptic fraction) were precipitated with 10 volumes of acetone at −20°C and recovered by centrifugation (20,000 g; 30 min, 4°C). The pellet containing insoluble postsynaptic density (PSD) was resuspended in 10 volumes solution buffer (1% Triton X-100, 20 μM Tris, pH 8.0, 4°C), incubated for 30 min, and centrifuged (40,000 g, 30 min, 4°C).

### GST-pull-down and co-immunoprecipitation assays

The methods used to produce the GST fusion proteins in bacteria have been previously reported (Anzai et al., 2002; Anzai et al., 2004). GST-pull-down assays were carried out using a GST Protein Interaction Pull-Down Kit (Pierce) according to the manufacturer’s instructions. Briefly, the bacterial lysate containing the fusion proteins was sonicated and centrifuged, and the supernatant was used for protein binding assays. In vitro translation was facilitated using a plasmid carrying full-length GluN2A, GluN2B or GluN1 (Addgene) that was transfected into HEK293T cells. The translated total proteins obtained from cells and the GST fusion proteins were incubated together with GST glutathione agarose resin. The protein complexes were then eluted by glutathione and analyzed by Western blotting using GST, GFP, GluN2A, GluN2B or GluN1 antibodies.

For co-immunoprecipitation assays, the extracts contained in the crude synaptosomal fraction obtained from adult mouse hippocampal tissues were prepared with anti-GluN2A, anti-GluN2B, anti-GluN1 or anti-FRMPD2 antibodies or IgG (rabbit, Abcam) (Control) at 4°C for 12 h before they were incubated with protein A/G agarose beads (Beyotime). The immunoprecipitated mixture was immunoblotted with GluN2A, GluN2B, GluN1 or FRMPD2 antibodies.

### Surface Plasmon Resonance (SPR) assays

SPR biosensor analysis was used to verify binding between peptides and the PDZ2 domain of FRMPD2 protein using a Biacore 8K instrument (GE Healthcare), as previously described(Zhang et al., 2014). Proteins were immobilized on S CM5 chips using amine-coupling chemistry. Binding experiments were performed in running buffer (10 mM PBS, pH 7, 0.05% P20) at 25°C with a flow rate of 30 μL/min. Gradient concentrations of PDZ2 proteins were diluted to 50 ug/ml in sodium acetate(10 mM, pH 5.5) and injected into the channel for 120 s, followed by disassociation for 120 s. RU values were collected at 10 Hz and all the experimental data were globally analyzed in a steady-state model within Biacore 8K Evaluation software.

### Study approval

All protocols of animal studies were approved by the Commission of Chongqing Medical University for the ethics of experiments on animals and conducted in accordance with international standards.

### Crystallization

GST-PDZ2 (amino acids 934-1027) was expressed in Escherichia coli, BL21(DE3) cells. Cells pellet was sonicated and centrifuged at 5000 rpm. The supernatant was subjected to GST column purification (10 mM Tris, 10 mM GSH, pH 7.4). Pooled fractions were dialyzed with thrombin and passed through a Q-HP column (20 mM Tris, 0-1 M NaCl, 5% glycerol, pH 8.0). Finally, protein fractions were further purified by Superdex 75 columns (20 mM Tris, 5% glycerol, 150 mM NaCl, 2 mM DTT pH 7.4) for crystallization.

Screen kits, including the JCSG-CORE-I-IV (Qiagen) and HR2-144 Index (Hampton), were used to perfrom crystallization screen with the hanging drop method at 18 °C. The protein (15 mg/ml) was mixed with an equivalent volume (1 μl) of screen buffer. The crystal was grown in 20% PEG3350 and 0.2 M sodium thiocyanate. A single crystal was transferred to the crystallization buffer containing an additional 20% glycerol and flash frozen with liquid nitrogen. The crystal diffracted well at BL19U (Shanghai Synchrotron Radiation Facility, SSRF, China). In all, images for 180° were collected. The data were processed by HKL3000 (Table 1). The crystal structure was refined by PHENIX and Coot. The structure was refined to 1.80 Å with R factor 19.76% and R free 24.76%.

### Statistical analysis

All averaged data are presented as the means ± SD, and all graphs were prepared using GraphPad Prism 4 software (La Jolla, CA). For independent-samples, Student’s t-test was used. Repeated measures ANOVA followed by post hoc t-tests was used to measure differences between two groups at multiple time points. All tests were two-sided. n indicates the number of cells/slices or independent experiments and was used to calculate the degree of freedom. All samples included in each experiment were analyzed in triplicate.

## Acknowledgements

This work was supported by grants from the National Natural Science Foundation of China (No. 81271445, 81471319, 81301109, and 81301110) and the National Clinical Key Specialty Construction Foundation of China.

## Author contributions

X. L. and XF. W. conceived and designed the experiments. HQ. Z. and X. L. performed the experiments. X. L. collected and analyzed the data. X. L. and X.T. wrote the manuscript. All authors contributed to preparation of the paper and approved the final contributions.

The authors have declared that no conflict of interest exists.

